# SUMOylation- and GAR1-dependent regulation of dyskerin nuclear and subnuclear localization

**DOI:** 10.1101/2020.09.02.280198

**Authors:** D.E. MacNeil, P. Lambert-Lanteigne, J. Qin, F. McManus, E. Bonneil, P. Thibault, C. Autexier

## Abstract

Dyskerin, a telomerase-associated protein and H/ACA ribonucleoprotein complex component plays an essential role in human telomerase assembly and activity. The nuclear and subnuclear compartmentalization of dyskerin and the H/ACA complex is an important though incompletely understood aspect of H/ACA ribonucleoprotein function. The posttranslational modification, SUMOylation, targets a wide variety of proteins, including numerous RNA-binding proteins, and most identified targets reported to date localize to the nucleus. Four SUMOylation sites were previously identified in the C-terminal Nuclear/Nucleolar Localization Signal (N/NoLS) of dyskerin, each located within one of two lysine-rich clusters. We found that a cytoplasmic localized C-terminal truncation variant of dyskerin lacking most of the C-terminal N/NoLS and both lysine-rich clusters represents an under-SUMOylated variant of dyskerin compared to wildtype dyskerin. We demonstrate that mimicking constitutive SUMOylation of dyskerin using a SUMO3-fusion construct can drive nuclear accumulation of this variant, and that the SUMO site K467 in this N/NoLS is particularly important for the subnuclear localization of dyskerin to the nucleolus in a mature H/ACA complex assembly- and SUMO-dependent manner. We also characterize a novel SUMO-interacting motif in the mature H/ACA complex component GAR1 that mediates the interaction between dyskerin and GAR1. Mislocalization of dyskerin, either in the cytoplasm or excluded from the nucleolus, disrupts dyskerin function and leads to reduced interaction of dyskerin with the telomerase RNA. These data indicate a role for dyskerin C-terminal N/NoLS SUMOylation in regulating the nuclear and subnuclear localization of dyskerin, which is essential for dyskerin function as both a telomerase-associated protein and as an H/ACA ribonucleoprotein involved in rRNA and snRNA biogenesis.

## Introduction

The H/ACA ribonucleoprotein (RNP) complex is responsible for pseudouridine synthesis at specific bases in ribosomal (r)RNA and small nuclear (sn)RNA in subnuclear compartments, specifically the nucleolus and Cajal bodies, respectively (1-5). The protein components of this complex at maturity are dyskerin (6-8), NOP10, NHP2 (9), and GAR1 (1, 10). The mature H/ACA complex assembles with noncoding (nc)RNA members of the H/ACA family, such as small nucleolar (sno)RNAs and small Cajal body specific (sca)RNAs that provide target pseudouridine synthesis specificity to dyskerin, the pseudouridine synthase of the H/ACA complex. The H/ACA motif is also a conserved biogenesis domain in telomerase RNAs of metazoans (11), including the human telomerase RNA (hTR) which relies on the H/ACA complex proteins for stability, processing, and function (12-16).

While hTR has no known target for guiding pseudouridine synthesis by dyskerin, the importance of the H/ACA complex in hTR biogenesis is demonstrated by mutations causing the premature aging disease and telomere syndrome dyskeratosis congenita (DC), with reported patient mutations in the genes encoding each protein component of the mature complex, excluding GAR1, as well as in the H/ACA biogenesis domain of hTR itself (17). Patients with DC have characteristic accelerated telomere shortening which leads to pathology in proliferative tissues, and results in bone marrow failure as the leading cause of mortality in this disease (18-21). In particular, DC patients with mutations disrupting the H/ACA complex components or H/ACA domain of hTR have reduced hTR accumulation which drives telomerase activity defects and accelerated telomere shortening (12). There have also been several reports of DC mutations in the H/ACA complex components affecting H/ACA RNA biogenesis beyond hTR (22, 23), and the essentiality of dyskerin and the H/ACA complex is likely due to its importance in rRNA and snRNA posttranscriptional modification. The *dkc1* gene encoding dyskerin is a core essential gene that is highly conserved, with phylogenetic roots in bacteria and archaea. Knockout of this gene is lethal in fungi (6), flies (24, 25), mice (26), and human cells (27, 28). Though X-linked dyskeratosis congenita (X-DC) is a commonly inherited form of the disease caused by mutations in *dkc1*, a complete deletion or loss of the gene has never been reported in X-DC, further demonstrating the essentiality of dyskerin.

The compartmentalization of dyskerin and the H/ACA complex is an important though incompletely understood aspect of H/ACA RNP function. Dyskerin has been reported to rely on two nuclear/nucleolar localization sequences (N/NoLSs) for complete nuclear import and retention, as well as for nucleolar accumulation (29). With the exception of GAR1, the H/ACA RNP components are present at sites of transcription of H/ACA RNAs in the nucleoplasm, along with the assembly factor NAF1 which is replaced by GAR1 upon complex maturation (30-32). Mature H/ACA complexes localize in the dense fibrillar component (DFC) of the nucleolus and in the Cajal bodies (7) where they guide posttranscriptional modification of rRNA and snRNA, respectively, dependent upon the H/ACA RNA with which the complex is assembled. The stepwise assembly of H/ACA RNPs has been proposed to play a role in localization of the complex to its sites of function (32). Although the mechanism governing subnuclear compartmentalization of the mature H/ACA complex remains incompletely characterized, it is likely to rely on regulation of miscibility with these discrete membrane-free regions of the nucleus. This has been recently demonstrated for other nucleolar proteins resident in the DFC such as fibrillarin, which relies on an intrinsically disordered glycine and arginine rich (GAR) domain and RNA interactions for miscibility with the DFC (33, 34).

The posttranslational modification SUMOylation has been demonstrated to affect nuclear and subnuclear localization of a number of protein targets, including resident proteins of the nucleolus (35-37). This modification involves conjugation of small ubiquitin-like modifier (SUMO) protein to lysine residues of target proteins in an E1 activating (SAE1/SAE2) and E2 conjugating (Ubc9) enzyme-dependent manner, often with the help of one of many E3 SUMO ligases, and promoted by a SUMOylation consensus motif in target proteins (ΨKXE/D – where Ψ is a hydrophobic residue and X is any residue) (38). While SUMOylation has been reported to regulate various functions of target proteins, a key aspect of SUMOylation is mediating protein-protein interactions between SUMO targets and proteins containing SUMO-interacting motifs (SIMs) which non-covalently bind SUMO (39-41). SUMOylation is a reversible modification, with several identified SUMO-specific proteases cleaving immediately after the C-terminal diglycine repeat in SUMO moieties, and therefore being responsible both for maturation of free SUMO and for removal of SUMO from target lysines (42-45). Typically, at steady state, only a small proportion of a SUMO target is conjugated to SUMO moieties. We previously demonstrated that dyskerin is a SUMOylation target of SUMO1 and SUMO2/3 isoforms, and that substituting either of two N-terminal X-DC-implicated lysine residues to arginine reduces the proportion of SUMOylated dyskerin in cells, leading to reductions in hTR, reduced telomerase activity, and accelerated telomere shortening (46). We have since shown that these two X-DC residues impact the dyskerin-hTR interaction, though the SUMO dependence of this interaction was not investigated (16).

Here we further investigate a regulatory role for SUMOylation of dyskerin. Using mutational analyses and SUMO-fusion constructs, we demonstrate that the C-terminal N/NoLS of dyskerin is a SUMO3 target, and that mimicking constitutive SUMOylation of a cytoplasmic truncation variant of dyskerin is sufficient to drive nuclear accumulation but not proper subnuclear localization of dyskerin. We also demonstrate that the nucleolar localization of dyskerin is mediated by the SUMO3 site K467 in this C-terminal N/NoLS, and that K467 is required for the interaction between dyskerin and GAR1 in a SUMO3-dependent manner, and novelly identify a SIM in GAR1 that is important for this interaction.

## Results

### The C-terminal nuclear/nucleolar localization sequence of dyskerin is a SUMOylation target

Many proteome-wide studies performed in human cell lines have identified dyskerin as a target of SUMOylation, both by SUMO1 and SUMO2/3 (47-55). Compiling the results of these studies, it is evident that dyskerin is a highly decorated target for SUMOylation, with 24 sites identified by mass spectrometry (MS) analyses (**Figure 1A**). For the purpose of this study, we focused on four SUMO2/3 sites in particular due to the placement of these lysines in the C-terminal N/NoLS (K467, K468, K498, and K507), which was previously reported to mediate efficient localization of dyskerin to the nucleus alone and in combination with an N-terminal N/NoLS (29). Importantly, truncation of the C-terminal N/NoLS by replacing K446 with a stop codon (X), and thus removal of all four SUMO3 sites and the lysine-rich (K-rich) clusters in which they are situated, substantially reduces the amount of SUMOylated dyskerin detectable by immunoblotting following Ni-NTA purification from HEK293 cells expressing FLAG-tagged dyskerin and 6xHis-SUMO3 (**Figure 1B**, wildtype vs. K446X). Indeed, while FLAG-tagged wildtype dyskerin co-localizes with the nucleolar marker fibrillarin in HEK293 cells assessed by immunofluorescence (IF), the FLAG-tagged K446X accumulates predominantly in the cytoplasm (**Figure 1C**, top and middle panels). Interestingly, mimicking constitutive SUMOylation of K446X by fusing a SUMO3 moiety to the N-terminus of this dyskerin variant allows for detection of high molecular weight products by Ni-NTA from HEK293 cells co-expressing FLAG-tagged SUMO3-K446X and 6xHis-SUMO3, indicating that this fusion protein is highly SUMOylated (**Figure 1B**). This SUMO3-fusion is also sufficient to drive the K446X truncation variant into the nucleus (**Figure 1C**, bottom two panels). However, the SUMO3-fusion variant remains excluded from the nucleolar compartment, suggesting that mimicking permanent SUMOylation of dyskerin disrupts proper subnuclear localization. This hypothesis is supported by our observation that fusion of SUMO3 to either the N-terminus or the C-terminus of FLAG-tagged wildtype dyskerin also leads to disrupted subnuclear localization (**Figure 1D**). These data suggest that the C-terminal N/NoLS of dyskerin regulates nuclear localization in a SUMO3-dependent manner, though the reversibility of SUMOylation after nuclear import is likely important for mediating proper subnuclear trafficking of dyskerin. This would be consistent with a previous proposal that “balanced SUMOylation levels” may be required for nucleolar regulation (56).

**Figure 1:**
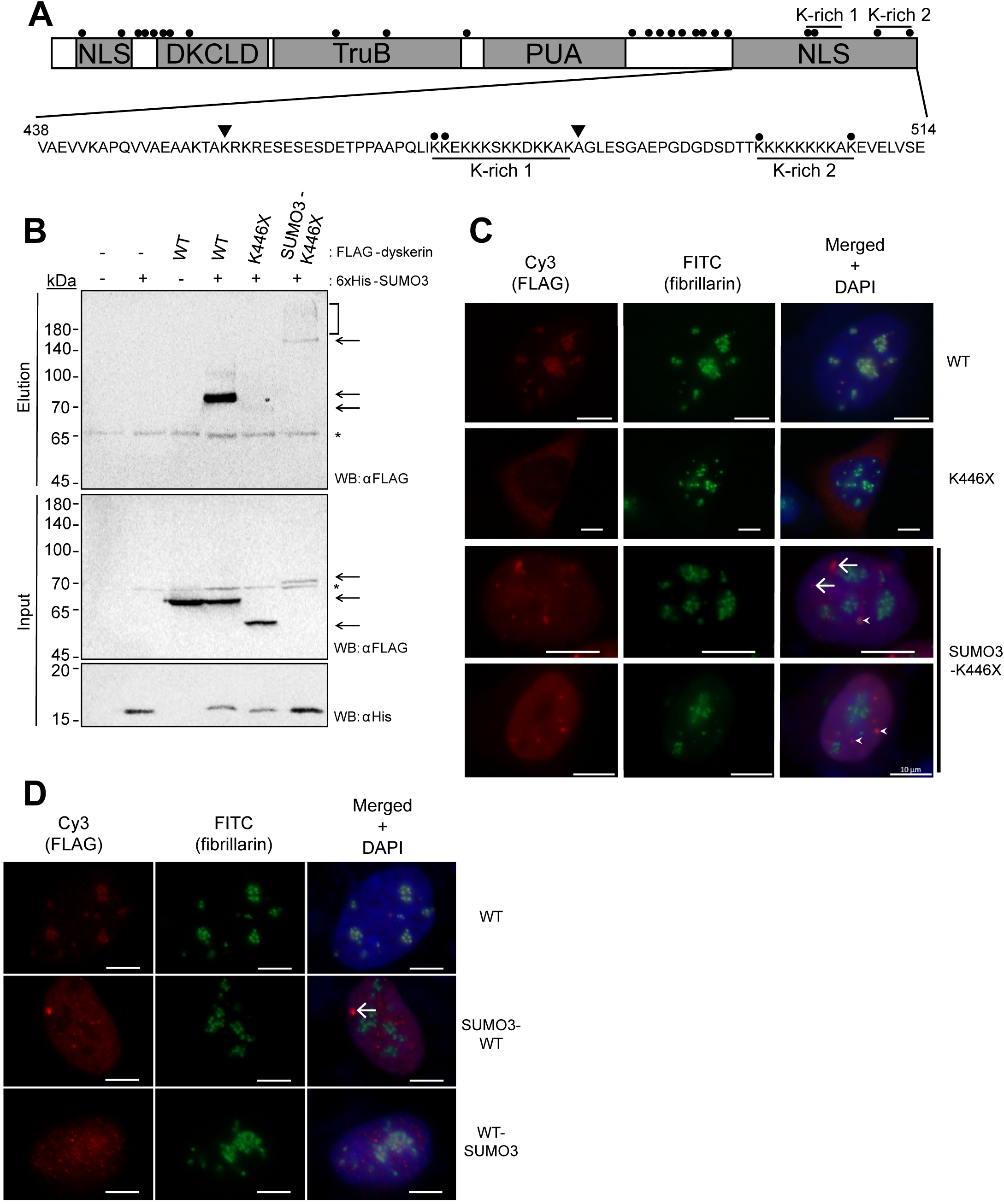
Residues in the C-terminal nuclear/nucleolar localization sequence of dyskerin are SUMO3 targets that govern nuclear accumulation. **a**. A linear schematic of human dyskerin domains. The amino acid range corresponding to the predicted C-terminal nuclear/nucleolar localization sequence (N/NoLS) (438-514) is denoted below the schematic, indicating the MS-identified SUMO3 sites in this region (K467, K468, K498, and K507) with solid black circles, and the two lysine (K)-rich clusters (K467-K480, and K498-K507) are underlined. MS-identified SUMO3 sites reported in proteome-wide studies cited in the text are indicated by solid black circles above the schematic. Residues A481 and K446 are indicated by arrowheads. **b**. FLAG-dyskerin (wildtype WT and dyskerin truncation variant K446X without or with N-terminal SUMO3 fusion) and 6xHis-SUMO3 were expressed in HEK293 cells. His-SUMO3 conjugates were purified using Ni-NTA agarose beads following lysis under denaturing conditions, and SUMOylated FLAG-dyskerin was assessed in the elution by immunoblotting with an anti-FLAG antibody. A fraction of each HEK293 cell pellet used for purification was kept prior to lysis (Input) for checking expression of FLAG-dyskerin and His-SUMO3 by immunoblotting. The K446X truncation runs at the expected lower molecular weight than WT dyskerin, while SUMO3-K446X runs at the expected higher molecular weight than WT due to the SUMO3 fusion. FLAG-dyskerin is indicated by arrows, while asterisks indicate non-specific antibody signal. The higher molecular weight smear observed in the SUMO3-K446X elution (indicated by the bracket to the right of the blot) indicates SUMOylated species of dyskerin. **c**. Representative images of the co-localization of FLAG-dyskerin (wildtype WT and dyskerin truncation variant K446X without or with N-terminal SUMO3 fusion – Cy3 shown in red) with nucleolar marker fibrillarin (FITC shown in green), as observed by indirect immunofluorescence. The nucleus is indicated by DAPI staining of nuclear DNA (in blue). **d**. Localization of FLAG-tagged WT, N-terminal SUMO3 fusion WT dyskerin (SUMO3-WT), and C-terminal SUMO3 fusion WT dyskerin (WT-SUMO3) was assessed by indirect immunofluorescence (Cy3 shown in red). Fibrillarin was used as a nucleolar marker (FITC shown in green), and the nucleus is indicated by DAPI staining of nuclear DNA (in blue). In **c**. and **d**. examples of nucleoplasmic FLAG-dyskerin foci that co-localize with nucleoplasmic fibrillarin foci are indicated by arrowheads, while arrows indicate nucleoplasmic FLAG-dyskerin foci that do not co-localize with nucleoplasmic fibrillarin foci. All scale bars indicate 10µm.

### Dyskerin nuclear and subnuclear localization is mediated by SUMOylation

The C-terminal N/NoLS of dyskerin contains two K-rich clusters, each of which contains two MS-identified SUMO3 target sites (**Figure 1A**). To elucidate which of these four SUMO3-sites, if any, may be mediating localization of dyskerin, and to reduce potential compensation for a loss of a single SUMOylation site by SUMO conjugation to neighbouring lysine residues, a stop codon was introduced at A481 in the FLAG-dyskerin construct, thus removing the entire second K-rich cluster while leaving the first intact. In contrast to the K446X variant, FLAG-tagged A481X efficiently localizes in the nucleus and the nucleolus, as observed by co-localization with fibrillarin assessed by IF (**Figure 2A**). This suggests that the second K-rich cluster, thus the SUMO3 sites within it are not critical regulators of dyskerin nuclear localization *per se*, and these data are consistent with previous localization analysis of a truncation variant at D493 (29). However, full length FLAG-tagged dyskerin in which the SUMO3 site K467 in the first K-rich cluster is substituted to an arginine (K467R) displays an apparent nucleolar exclusion/nucleoplasmic accumulation phenotype when assessed by IF (**Figure 2A**). This is in contrast to the K468R variant that localizes comparably to wildtype dyskerin (**Figure 2A**). Substituting both of these lysines to arginine (K467/468R) leads to a localization phenotype similar to the single K467R substitution variant (**Figure 2A**). FLAG-positive cells were scored based on localization phenotype as a percentage of FLAG-positive cells counted, and localization of each FLAG-tagged dyskerin (wildtype or variant) was assessed from three independent experimental replicates (**Figure 2B, C**). Importantly, while the major localization phenotype of the K446X truncation variant is cytoplasmic, FLAG-signal in cells expressing this variant was also observed in both the cytoplasm and nucleolar fraction concomitantly (**Figure 2B**, blue bar). This is consistent with previous time course experiments demonstrating through microinjection of EGFP-tagged K446X into cells that this truncation impairs but does not entirely prevent nuclear and subnuclear localization of dyskerin (29). While still able to localize within the nucleus and to the nucleolus, the truncation variant of dyskerin at A481, A481X, does display an increase in concomitant nucleoplasmic and nucleolar localization of dyskerin compared to wildtype, suggesting that this truncation modestly affects localization of dyskerin, albeit to a lesser extent than K446X or K467R (**Figure 2B**, brown bar). Indeed, K467R has a substantial reduction in nucleolar and corresponding increase in nucleoplasmic localization compared to wildtype dyskerin, as assessed by exclusion from co-localization with fibrillarin signal, but co-localization with DAPI signal (**Figure 2B**, red and black bars, respectively). In contrast, no substantial differences in localization were observed between K468R and wildtype (**Figure 2B**). The double substitution variant K467/468R does not differ in localization compared to the single K467R variant, and thus has a reduction in nucleolar and increase in nucleoplasmic localization compared to wildtype dyskerin (**Figure 2B**). As previously shown (**Figure 1C**), fusing K446X to SUMO3 is sufficient to drive this truncation into the nucleus, but this fusion variant has reduced nucleolar localization compared to wildtype dyskerin (**Figure 2B**). However, SUMO3-fusion of dyskerin differs from K467R in nucleolar exclusion, as SUMO3-K446X and SUMO3-wildtype dyskerin form distinct puncta in the nucleoplasm while K467R localization in the nucleoplasm is diffuse (**Figure 1C and D, Figure 2A**). Some of these puncta may represent CB’s given their occasional overlap with fibrillarin puncta outside of nucleolar clusters (**Figure 1C** arrowheads), but are most likely nucleoplasmic aggregates driven by the permanent nature of the SUMO3-fusion (**Figure 1C and D** arrows). Importantly, neither the FLAG-tag nor another tag (eGFP) disrupt localization of wildtype dyskerin, as eGFP-tagged wildtype dyskerin (**Figure 2D**) and endogenous dyskerin examined by IF (**Figure 2E**) display comparable localization patterns to exogenously expressed FLAG-tagged wildtype dyskerin. Taken together, these data tell us that in addition to SUMO3 mediating nuclear localization of the K446X truncation of dyskerin, the SUMO3 site K467 plays an important regulatory role for the nucleolar localization of dyskerin.

**Figure 2:**
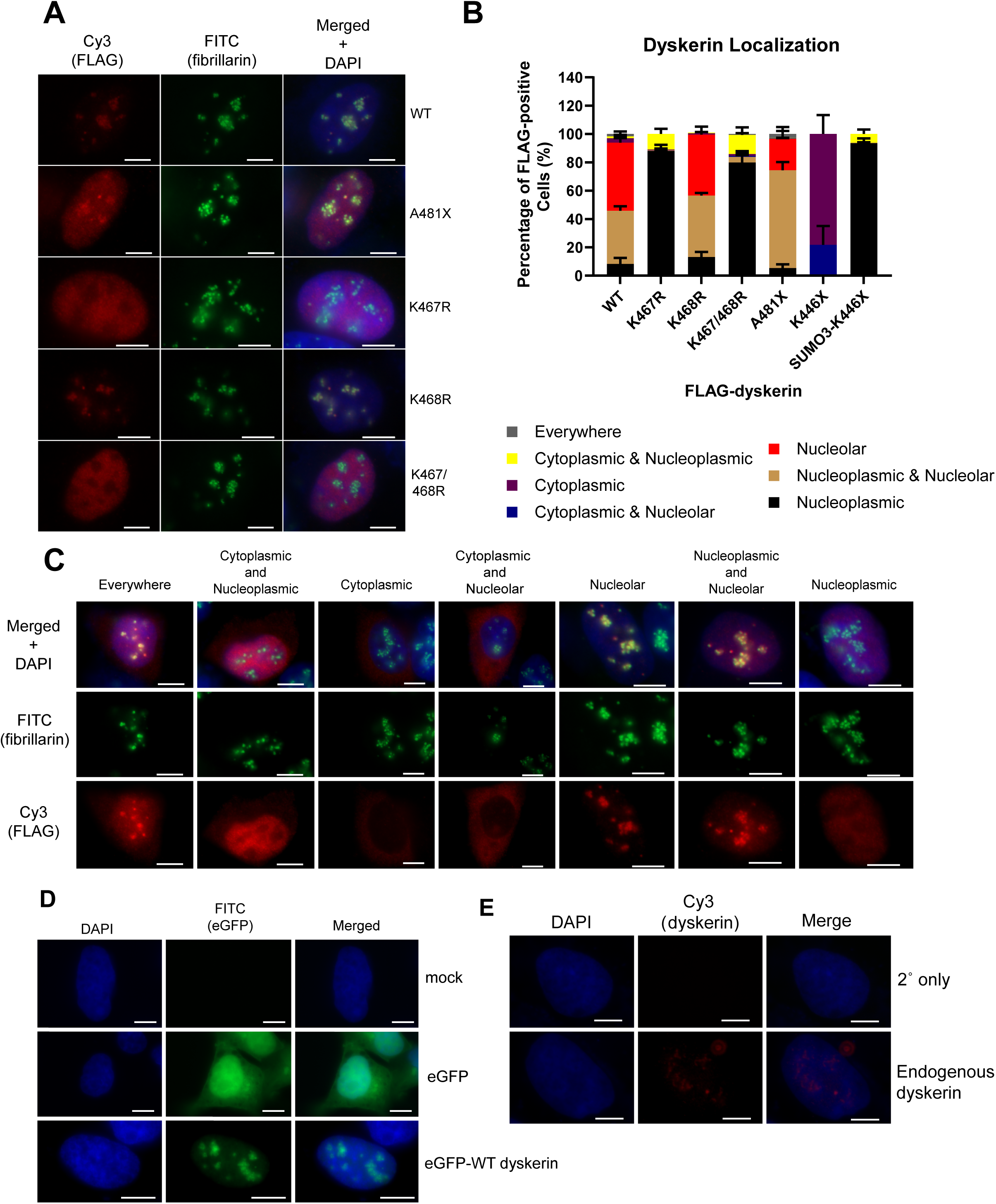
Nuclear and subnuclear localization of dyskerin is mediated by SUMO3 sites in the C-terminal nuclear/nucleolar localization sequence. **a**. FLAG-dyskerin was transiently expressed in HEK293 cells, and localization was assessed in fixed cells by indirect immunofluorescence. Representative images of the most prevalent localization phenotype of FLAG-dyskerin (wildtype WT, dyskerin truncation variant A481X, and substitution variants K467R, K468R, and double K467/468R – Cy3 in red) and the nucleolar marker fibrillarin (FITC in green) are shown. Nucleolar localization is represented for WT and K468R, concomitant nucleoplasmic & nucleolar is represented by A481X, and nucleoplasmic localization is represented by K467R and K467/468R. The nucleus is indicated by DAPI staining of nuclear DNA (in blue) **b**. Quantification of localization phenotype scoring for FLAG-dyskerin WT and localization variants as a percentage of FLAG-positive HEK293 cells is indicated (≥50 cells per condition were counted, in experiment replicate n=3). **c**. Examples of how localization phenotype was scored are shown. Nucleolar localization was determined by co-localization with clustered fibrillarin signal, nucleoplasmic localization was determined by co-localization with DAPI outside of the clustered fibrillarin signal, and cytoplasmic localization was determined by concentrated signal outside of and surrounding DAPI. **d**. Localization of eGFP and eGFP-tagged WT dyskerin (in green) was assessed in fixed HEK293 cells following transient transfection, using a FITC filter. **e**. Localization of endogenous dyskerin (Cy3, in red) was assessed by IF in fixed HEK293 cells. As a negative control, fixed HEK293 cells were assessed by IF using only secondary Cy3-conjugated antibody. The nucleus is indicated by DAPI staining of nuclear DNA (in blue). These are representative images.All scale bars indicate 10µm.

### Nuclear and subnuclear localization of dyskerin affects mature H/ACA RNP assembly

As a functional readout for H/ACA complex assembly and localization, co-immunoprecipitation (co-IP) of FLAG-tagged dyskerin and interacting components was performed from HEK293 cell lysate. Following FLAG-IP, interactions of FLAG-tagged dyskerin wildtype and N/NoLS variants were assessed by immunoblotting for endogenous H/ACA RNP assembly factors and components. Comparable to wildtype dyskerin, all FLAG-tagged N/NoLS variants were able to interact with the pre-H/ACA RNP component NAF1 and the pre- and mature H/ACA RNP components NOP10 and NHP2 (**Figure 3A**). Strikingly, the N/NoLS variants with nucleolar exclusion phenotypes (K467R and K467/468R) were unable to interact with the mature H/ACA RNP component GAR1 (**Figure 3B**). Importantly, disruption of either nuclear or subnuclear localization of dyskerin leads to impaired hTR-dyskerin interaction as measured by qPCR following RNA extraction and reverse transcription from IP fractions; neither FLAG-tagged K467R nor K446X interact with hTR relative to wildtype dyskerin (**Figure 3C**). This is in contrast to the N/NoLS variants with little to no localization defects, K468R and A481X which do not display defective interactions with hTR relative to wildtype dyskerin (**Figure 3C**). These data indicate that proper localization of dyskerin is tied to H/ACA RNP complex assembly, connect GAR1-dyskerin interaction defects to the nucleolar exclusion of dyskerin, and demonstrate that improper dyskerin localization disrupts the ability of dyskerin to interact with H/ACA RNAs like hTR.

**Figure 3:**
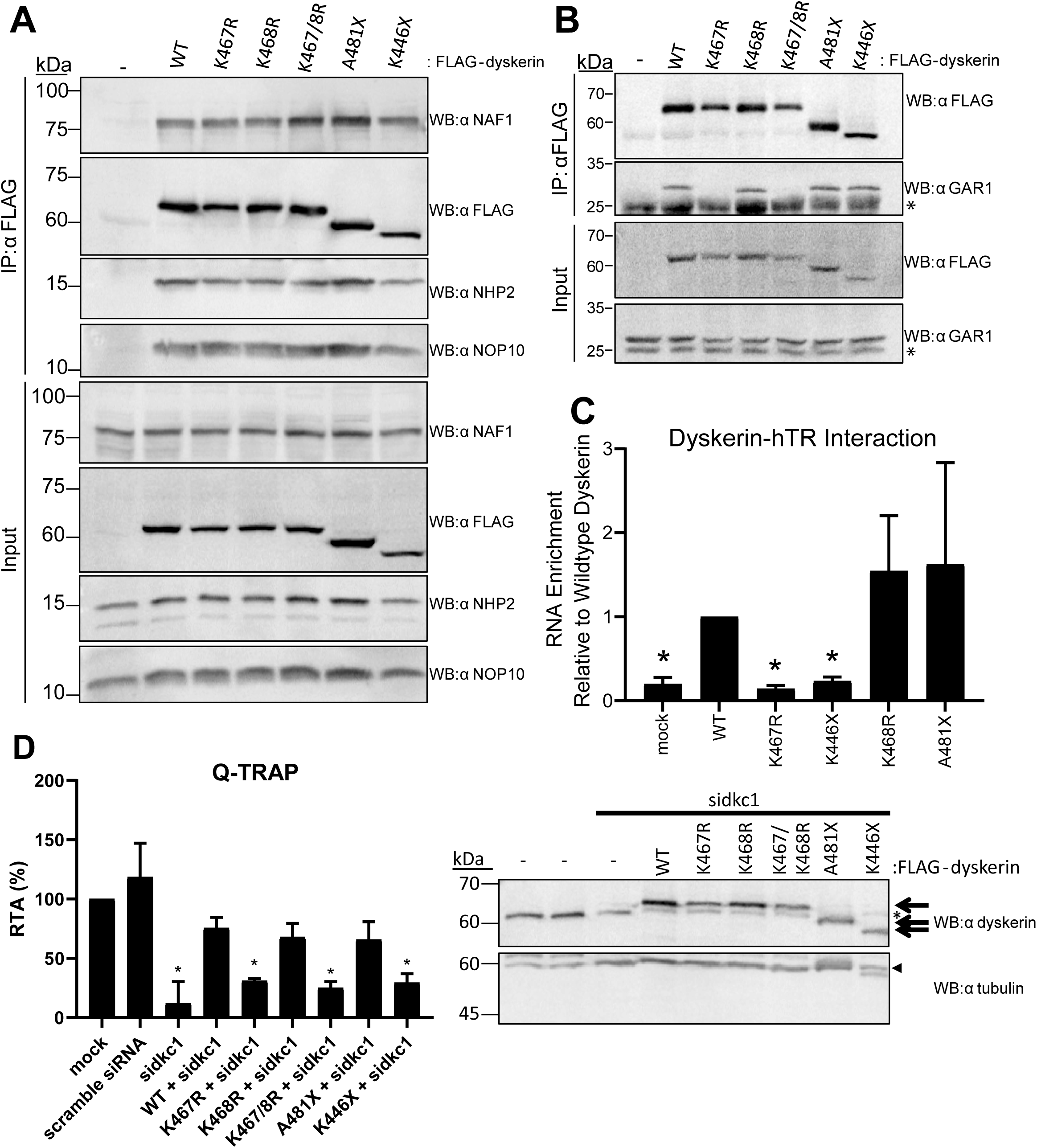
Dyskerin nuclear and nucleolar localization is linked to mature H/ACA complex assembly and function. Interactions of FLAG-dyskerin WT and localization variants with endogenous pre- and mature H/ACA ribonucleoprotein complex components were assessed by co-immunoprecipitation (IP) from HEK293 cell lysates. Assembly of the **a**. H/ACA pre-RNP complex involving NAF1, NHP2, and NOP10 was investigated by immunoblotting for the endogenous H/ACA pre-RNP components and FLAG-dyskerin proteins following IP. **b**. Interaction of dyskerin with the mature H/ACA complex component GAR1 was examined following IP by immunoblotting for endogenous GAR1 and FLAG-dyskerin. Localization variants that are excluded from the nucleolus (K467R and K467/468R) do not interact with GAR1. Immunoblotting targets are indicated to the right of each panel as “WB: α target”, and a list of antibodies can be found in the materials and methods section. A non-specific band revealed by the anti-GAR1 antibody at 25kDa is indicated by an asterisk. Each co-IP and immunoblotting was performed in experimental replicate a minimum of n=2, representative blots are shown. **c**. Dyskerin-hTR interactions were assessed by IP of FLAG-tagged dyskerin followed by RNA extraction and qPCR. Relative to wildtype IP fractions, dyskerin variants with substantial localization defects (K467R and K446X) display significantly reduced enrichment of hTR following IP. HEK293 cells lacking FLAG-tagged dyskerin (indicated as mock) were used as a negative control for RNA binding to the FLAG antibody and/or Protein G Sepharose. Mock cells were subject to the same IP protocol detailed for fractions containing FLAG-tagged dyskerin. These data represent experimental replicates of n=3. Statistically significant reductions in enrichment relative to wildtype are indicated by * (P value < 0.01). Error bars represent SEM.**d**. HEK293 cells without or with stable expression of FLAG-dyskerin (WT or N/NoLS variants) were depleted of endogenous dyskerin using siRNA, and confirmed by immunoblotting against endogenous dyskerin. The same membrane was first probed using an anti-dyskerin antibody, followed by an anti-tubulin antibody as a loading control. Arrows indicate FLAG-tagged dyskerin, the asterisk indicates endogenous dyskerin, and the arrowhead indicated tubulin. Q-TRAP was performed using cell lysate. Statistically significant reductions in relative telomerase activity (RTA) compared to mock (untreated) HEK293 cells are indicated by * (P value < 0.0001), and this was repeated in experimental triplicate. Error bars represent SEM.

### Defects in dyskerin localization affect telomerase activity

We asked whether H/ACA complex assembly defects disrupted dyskerin function in the context of telomerase activity and H/ACA RNA biogenesis. In order to assess telomerase activity, endogenous dyskerin was depleted via siRNA targeting the 3’ UTR of dyskerin in HEK293 cells with or without stable expression of FLAG-tagged dyskerin wildtype or N/NoLS variants. After 72h of depletion, telomerase activity was measured in the cell lysate using Q-TRAP. HEK293 cells depleted of endogenous dyskerin have significantly reduced telomerase activity compared to untreated (mock) cells, cells treated with a scramble siRNA, and cells with stable exogenous expression of FLAG-tagged wildtype dyskerin that are depleted of endogenous dyskerin **(Figure 3D)**. Additionally, HEK293 cells with stable expression of FLAG-tagged dyskerin localization variants that have defective localization (K467R, K467/8R, and K446X) display significantly reduced telomerase activity following depletion of endogenous dyskerin. In contrast, cells with stable expression of FLAG-tagged dyskerin localization variants that are competent for nuclear and nucleolar localization (K468R and A481X) display telomerase activity similar to cells with stable expression of FLAG-tagged wildtype dyskerin following depletion of endogenous dyskerin.

### GAR1 interaction with dyskerin mediates nucleolar localization in a SUMO-dependent manner

While the dyskerin variants that are excluded from the nucleolus do not interact with GAR1, the mainly cytoplasmic K446X truncation which is competent for nucleolar localization is capable of interacting with endogenous GAR1 (**Figure 2B, Figure 3B**). Due to this observation that nucleolar localization of dyskerin is connected to the dyskerin-GAR1 interaction and the SUMO3 site K467, we asked whether the interaction between GAR1 and dyskerin may be SUMO3-mediated, and whether this interaction is responsible for mediating dyskerin nucleolar localization. To test this, we performed FLAG co-IPs from HEK293 cells expressing FLAG-tagged dyskerin fused to SUMO3 at the C-terminus. Interestingly, fusing SUMO3 to the C-terminus of the K467R variant (**Figure 4A**) is able to rescue the robust GAR1 interaction defect of K467R (**Figure 3B**), and wildtype dyskerin with C-terminal SUMO3 fusion is also able to interact with GAR1 comparably to wildtype dyskerin alone (**Figure 4A**). However, the subnuclear localization of both of these SUMO3-fusions does differ from that of wildtype dyskerin, as assessed by IF, consistent with a predicted requirement for SUMOylation reversibility for proper regulation of dyskerin subnuclear localization. Importantly, compared to fusion of SUMO3 to the N-terminus of K446X or the non-fusion K467R variant, these C-terminal SUMO3 fusions display more co-localization with fibrillarin in the nucleolar compartment, though less frequent exclusive nucleolar localization of these fusions is observed compared to wildtype dyskerin (**Figure 2B, Figure 4B**). This suggests that the nucleolar localization of K467R, as well as interaction between GAR1 and dyskerin may indeed be SUMO3-dependent. To further elucidate a potential SUMO3-mediated GAR1-dyskerin interaction, and using the prediction software GPS-SUMO 4.0, we identified a single predicted SIM within GAR1 at residues 70-74 (70-VVLLG-74) proximal to the previously predicted dyskerin-GAR1 interface (**Figure 4C**). In order to assess whether this predicted SIM could mediate the interaction between GAR1 and dyskerin, we substituted each residue in the predicted SIM to alanine in a 3xFLAG-tagged GAR1 construct (annotated as 5A), and assessed the interaction of 3xFLAG-tagged GAR1 with endogenous dyskerin. Wildtype 3xFLAG-tagged GAR1 is able to interact with endogenous dyskerin, as assessed by FLAG co-IP from HEK293 cells expressing 3xFLAG-GAR1, however GAR1 5A displays a reduced interaction with endogenous dyskerin (**Figure 4D**). These data demonstrate that GAR1 contains a SIM which mediates the efficient interaction between dyskerin and GAR1 in a SUMO3-dependent manner, relying on the SUMO3 site K467 in the C-terminal N/NoLS of dyskerin, and that this interaction with GAR1 governs the localization of dyskerin in the nucleolus.

**Figure 4:**
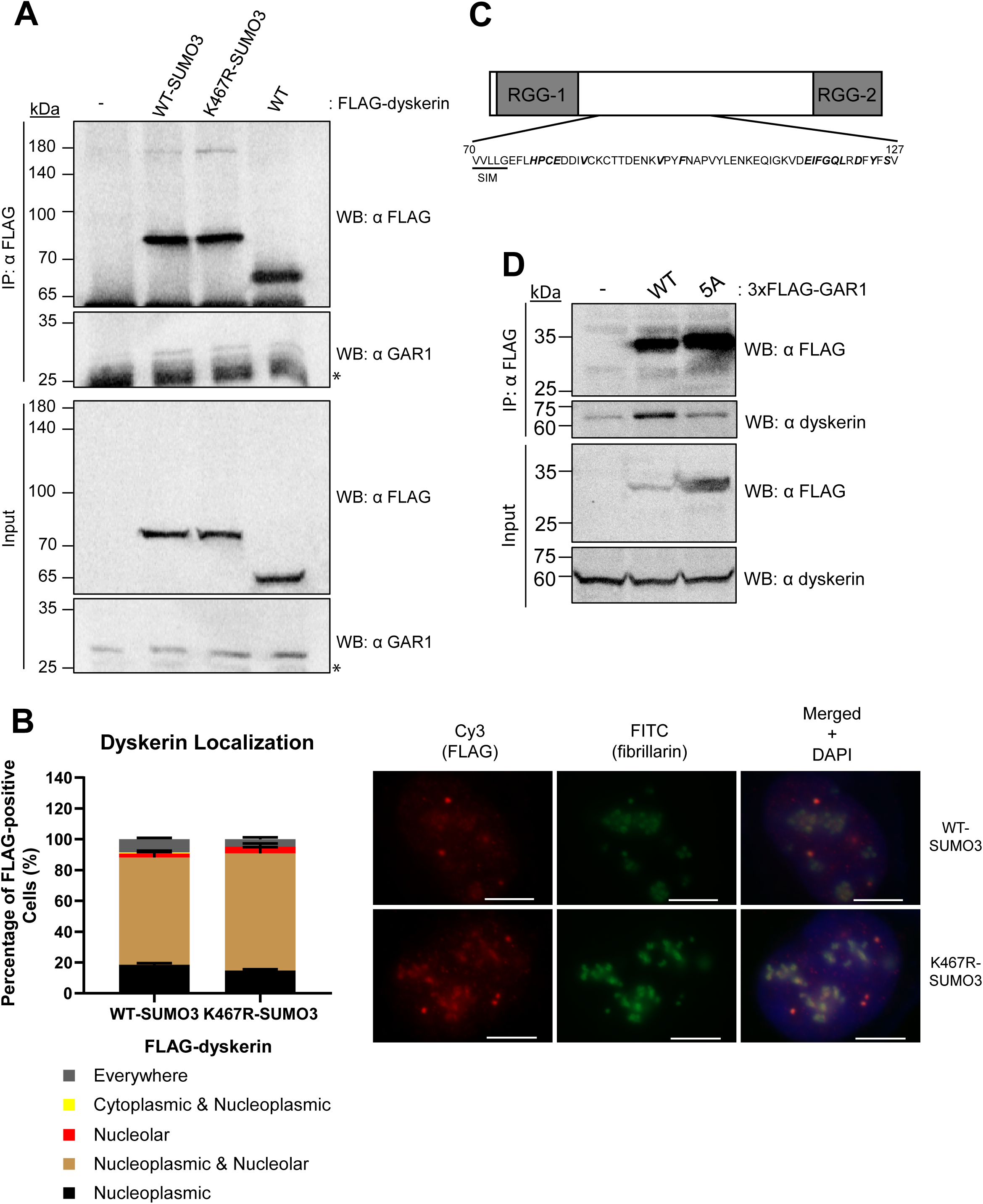
Efficient interaction between dyskerin and GAR1 is mediated by SUMO3. **a**. Following transient transfection of FLAG-tagged dyskerin in HEK293 cells and co-immunoprecipitation (IP) from cell lysates using FLAG antibody, the interaction between endogenous GAR1 and wildtype (WT)-SUMO3 fusion, K467R-SUMO3 fusion, or WT dyskerin was assessed by immunoblotting. SUMO3 fusion variants run at the expected higher molecular weight than WT dyskerin alone due to the SUMO3 moiety. A non-specific band revealed by the anti-GAR1 antibody at 25kDa is indicated by an asterisk. **b**. C-terminal SUMO3 fusions of FLAG-dyskerin wildtype and K467R variant were transiently expressed in HEK293 cells, and localization was assessed in fixed cells by indirect immunofluorescence. Representative images of the most prevalent localization phenotype of FLAG-dyskerin (Cy3 in red) and the nucleolar marker fibrillarin (FITC in green) are shown. The nucleus is indicated by DAPI staining of nuclear DNA (in blue). Quantification of localization phenotype scoring as a percentage of FLAG-positive HEK293 cells is indicated (≥50 cells per condition were counted, in experiment replicate n=3), and scale bars indicate 10µm. **c**. A linear schematic of human GAR1, with glycine and arginine rich domains indicated (RGG-1 and RGG-2). The predicted SIM (70-VVLLG-74) is indicated, and residues expected to physically interface with dyskerin based on previous homologue structural studies in yeast are bolded and italicized. **d**. Substitution of all five predicted SIM residues to alanine (GAR1 5A) impairs the interaction between GAR1 and dyskerin, as demonstrated by co-IP of 3xFLAG-GAR1 and endogenous dyskerin from HEK293 cells transiently expressing 3xFLAG-GAR1. Interaction between endogenous dyskerin and WT or 5A GAR1 was assessed by immunoblotting following FLAG IP from cell lysates. Immunoblotting targets are indicated to the right of each panel as “WB: α target”, and a list of antibodies can be found in the materials and methods section. Each co-IP and immunoblotting was performed in experimental replicate a minimum of n=2, representative blots are shown.

## Discussion

Dyskerin and the H/ACA RNP complex play essential roles in H/ACA RNA biogenesis, posttranscriptional modification of rRNA and snRNA, and in human telomerase assembly and activity. The ability of dyskerin to carry out its various functions relies heavily on its nuclear and subnuclear compartmentalization, where it assembles with H/ACA RNA and localizes to sites of function, including the nucleolus where pseudouridine synthesis occurs on rRNA. In this study we further demonstrate the interconnectedness of dyskerin localization, H/ACA RNP assembly, and complex function. More specifically, we demonstrate that efficient nuclear localization of dyskerin, driven by the K-rich C-terminal N/NoLS, as well as mature H/ACA complex assembly and nucleolar localization mediated by K467 in this N/NoLS are crucial for dyskerin assembly with H/ACA RNA like hTR. Furthermore, we demonstrate that the localization of dyskerin can be mediated by SUMOylation sites in the C-terminal N/NoLS.

A previous study of dyskerin nuclear localization characterized two N/NoLS regions, one in the N-terminus (amino acids 11-20) and one in the C-terminus (amino acids 446-514) (29). This foundational study reported that removing or mutating the N-terminal region alone did not disrupt localization of dyskerin, whereas removal of the C-terminal region alone drastically impeded nuclear localization, and combinatorial removal of both regions abolished nuclear localization altogether. As such, we focused on this C-terminal N/NoLS region as the primary driver of dyskerin nuclear localization. Strikingly, we found that efficient localization of dyskerin to the nucleus, while impaired by truncation of the C-terminal N/NoLS at K446, can be driven by mimicking SUMOylation through fusing dyskerin to a SUMO3 moiety. This suggests that the loss of SUMOylation sites from this truncation variant may be responsible for inefficient dyskerin nuclear import and/or retention. Given the multitude of dyskerin SUMOylation sites identified by MS which fall outside of the C-terminal N/NoLS of dyskerin, it is also likely that dyskerin SUMOylation takes place within the nucleus for some sites following nuclear import. Establishing which SUMOylation sites in particular govern nuclear localization requires further investigation, as pinpointing the MS-identified SUMOylation sites in this region responsible for nuclear localization was not evident by removal or substitution of K467, K468, K498, or K507. However, this mutational analysis instead revealed that K467 plays an important regulatory role in subnuclear localization of dyskerin to the nucleoli. Importantly, in our study and in the previous work by Heiss *et al*., full truncation of the C-terminal region does not prevent nucleolar localization of dyskerin *per se* (29). As such, we postulate that the C-terminal N/NoLS may govern several aspects of stepwise dyskerin localization (nuclear import, nucleoplasmic assembly with H/ACA RNA, and nucleolar miscibility) through regulated conformational changes. More specifically, we speculate that a conformational change of this C-terminal region governed by SUMOylation at K467 may be responsible for licensing dyskerin nucleolar localization. However, the absence of this region as a whole allows for dyskerin nucleolar localization in the absence of K467 SUMOylation, albeit in the context of inefficient nuclear localization, because no conformational change is required for the K446X truncation variant. This would be consistent with reports that full length dyskerin and dyskerin homologues are difficult to purify *in vitro* due to insolubility issues which can be resolved by removal of this C-terminal region (32, 57), and also in agreement with a lack of reported structure of this functionally required tail due to its apparent intrinsic low complexity (6, 58, 59). It also seems likely that a conformational change in dyskerin may be responsible for regulating the exchange of GAR1 for NAF1 upon H/ACA complex maturation, though this needs further investigation.

Meanwhile, subnuclear localization of dyskerin-SUMO3 fusion proteins, variant or wildtype, was observed to differ from wildtype dyskerin alone. We postulate that the constitutive nature of this SUMOylation mimic disrupts nucleolar localization due to the inability of deSUMOylating proteases to reverse this imitated posttranslational modification. This proposal is based on not only the abundance of nucleolar SUMO-targets and SUMOylation machinery involvement in nucleolar integrity (60-64), but also on the nucleolar localization of SUMO-specific proteases (SENP3 and SENP5) involved in deconjugation of SUMO2/3 from target proteins (65). SENP3 in particular has been demonstrated to interact with the nucleolar resident protein nucleophosmin (NPM1), the 60S maturation factors PELP1, TEX10, WDR18, and Las1L, and is capable of deSUMOylating NPM1, PELP1, and Las1L (66-68). Consistent with the hypothesis that SUMO removal may regulate nucleolar localization of SUMO-target proteins, depletion of SENP3, and thus reduction of nucleolar deSUMOylation, has been reported to lead to nucleolar release of the PELP1-TEX10-WDR18 complex (68). Furthermore, in yeast the nucleolar SUMO-specific protease Ulp2 has been demonstrated to reverse SUMOylation of rDNA-bound SUMO-targets, and engineered increased SUMOylation by depletion of Ulp2 leads to a reduction of several nucleolar proteins bound to rDNA (69, 70). Intriguingly, NPM1 is responsible for localization of SENP3 to the nucleolus (71). NPM1 is a resident protein of the outer-most nucleolar component where ribosomal subunit maturation takes place, the granular component (GC), and as such would make a good candidate for gatekeeping localization of nucleolar proteins in a SUMO removal-dependent manner. This remains uninvestigated but would also fit into models of phase-mediated nucleolar compartmentalization (33, 72), as discussed in greater detail below.

Here we also report that the efficient interaction between dyskerin and GAR1 is mediated through a newly characterized SIM in GAR1 (amino acids 70-VVLLG-74). SIMs are typically short hydrophobic stretches of residues that can form an extended β-strand backbone, which then non-covalently interacts with SUMO moieties to foster stronger or more frequent SUMO-mediated protein-protein interactions (38). It is important to note that this predicted motif is not well conserved in lower eukaryotes or archaea (58). We found that substituting all five of these GAR1 residues to alanine impairs the interaction of GAR1 with endogenous dyskerin, indicating that an efficient interaction between GAR1 and dyskerin relies on this SIM, which is proximal to but does not overlap with any of the residues structurally identified previously to mediate the interaction between these two proteins in yeast and archaea (58, 73). Anecdotally, this SUMO-mediated interaction between GAR1 and dyskerin may also offer some explanation for the reported difficulty of *in vitro* reconstitution of H/ACA complexes using full length proteins, and indeed the GAR1-dyskerin interface that has been identified structurally using homologues from other organisms does not account for the C-terminal N/NoLS of human dyskerin as this region was absent from the dyskerin homologues used for crystallization (58, 59, 74, 75). These structural data also indicate that the GAR1-dyskerin interaction does take place in the absence of SUMOylation and without the dyskerin C-terminal N/NoLS *in vitro*, indicating that while this GAR1 SIM contributes to the efficient interaction between dyskerin and GAR1 in a cellular context, this SIM is not required *per se*. We also observed that substituting the dyskerin SUMO3 site K467 to arginine abolishes the interaction between GAR1 and dyskerin, and that this GAR1 interaction defect of the K467R variant can be rescued by fusing K467R to SUMO3. It is not known if K467 directly interfaces with GAR1, due to the absence of data on this C-terminal region of dyskerin from structural studies. However, the observation that fusion of K467R to a SUMO3 moiety can recover the ability of this variant to interact with GAR1 strongly implies that SUMOylation of K467 mediates the efficient interaction between GAR1 and dyskerin.

Finally, we postulate that the SUMO-mediated interaction between GAR1 and dyskerin is required for dyskerin localization to the nucleolus. Along with the data we present here, this hypothesis is rooted in recent analyses of the nucleolar resident protein fibrillarin. Fibrillarin is a small nucleolar RNP counterpart to dyskerin responsible for the 2′O-methylation posttranscriptional modification of rRNA in the DFC, guided by C/D box snoRNA rather than H/ACA box snoRNA (76-80). Several studies have demonstrated that localization of fibrillarin to the DFC is mediated by an intrinsically disordered GAR domain, as well as by interactions with nascent pre-rRNA as the RNA is sorted radially from its site of transcription through the three nucleolar components, of which the DFC is the centre (33, 34). These studies and others have shown that the nucleolus represents a complex membrane-free compartment with three distinctly liquid-liquid phase separated components, which as a whole are phase separated from the surrounding nucleoplasm (33, 72, 81-83). This context is important to bear in mind when considering dynamic localization of resident nucleolar proteins in and out of these separated phases. The regulated miscibility of fibrillarin with the DFC relies on its GAR domain and protein-RNA interactions. As such, we propose that dyskerin miscibility with the DFC relies on its interaction with GAR1, not only through acting as a GAR domain for dyskerin and the entire H/ACA complex *in trans*, but also by providing high H/ACA complex-to-guide RNA affinity which facilitates accurate H/ACA complex placement on target RNA, like rRNA in the nucleolus (84, 85). This hypothesis is supported by our observations that 1) the K467R dyskerin variant is unable to interact with GAR1 and the H/ACA box RNA hTR; 2) this K467R variant is unable to co-localize with fibrillarin in the nucleolus; and 3) improving the interaction between the K467R variant and GAR1 by fusing K467R to SUMO3 also allows for partial co-localization of the K467R variant with fibrillarin in the nucleolus. Furthermore, the ability of the nucleolar-miscible K446X truncation to fully assemble with H/ACA pre-and mature RNP components, including interacting with GAR1 also lends support to this hypothesis. We also speculate that the lack of GAR domains in the archaeal homologues of GAR1 and fibrillarin provides evolutionary support for the notion that GAR domains mediate membrane-free compartmentalization of these complexes in eukaryotes, as archaea lack nuclear compartmentalization altogether and would have no need for GAR domain-mediated nucleolar miscibility of the otherwise evolutionarily conserved H/ACA or C/D RNP complexes (86). Further confirmation of the phase dynamics of human dyskerin with or without GAR1 is needed to elucidate this hypothesis.

## Methods

### Plasmids, Cell Culture, and Transfections

The plasmid pcDNA3.1-FLAG-dyskerin^WT^ from the lab of Dr. François Dragon was used to generate point mutations or truncations via site directed mutagenesis, as previously described (16, 46). Specifically, primers (**Table S1**) were designed to generate K467R, K468R, K467/468R, A481X, and K446X. For expression of 3xFLAG-GAR1 in human cells, the pcDNA3.1 3xF-GAR1 plasmid was purchased from Addgene (#126873), and the predicted SIM 70-VVLLG-74 was substituted to 70-AAAAA-74 by site directed mutagenesis. The construct pcDNA3.1-6xHis-SUMO3 was obtained from Dr. Frédérick Antoine Mallette (Université de Montréal). The plasmid encoding eGFP-tagged dyskerin (pmEGFP-C1-DKC1) was a gift from the lab of Dr. Ling-Ling Chen (Shanghai Institute of Biochemistry and Cell Biology) (34). All constructs underwent Sanger DNA sequencing at Génome Québec CES.

Human embryonic kidney (HEK293) cells were maintained in Dulbecco’s Modification Eagle’s Medium DMEM (Wisent) supplemented with 10% fetal bovine serum FBS (Wisent), and Antibiotic-Antimycotic (Gibco), at 37°C 5% CO_2_. Polyclonal FLAG-dyskerin stable cells were maintained under selective pressure in G418 (750µg/ml). Transfection of pmEGFP-C1-DKC1, pcDNA3.1-FLAG-dyskerin constructs, pcDNA3.1-6xHis-SUMO3, and/or pcDNA3.1 3xFLAG-GAR1 was performed using Lipofectamine 2000 Transfection Reagent (Invitrogen) according to the reagent protocol. Prior to transfection, media was changed to DMEM with 10% FBS and lacking Antibiotic-Antimycotic, and 5 hours after transfection the media was replaced with DMEM containing both FBS and Antibiotic-Antimycotic.

Transfection of siRNA was performed with Lipofectamine RNAiMAX Transfection Reagent (Invitrogen) according to the manufacturer’s user protocol. siRNA targeting the 3’ UTR (24 nM, 72h treatment, sidkc1) was used for depletion of endogenous dyskerin (**Table S1**). The siRNA sequence targeting the 3’ UTR was previously described (87). A mock transfection (no siRNA) and transfection of a scramble siRNA were used as negative controls in each experiment. siRNAs were ordered through ThermoFisher Scientific.

### SUMO-interaction Motif Prediction

The GPS-SUMO 4.0 prediction tool was used to predict possible SUMO-interacting motifs in GAR1. The coding amino acid sequence for isoform 1 of GAR1 was obtained in FASTA format through Uniprot (identifier Q9NY12-1). The “SUMO Interaction Threshold” was set to “Medium”. The SUMO Interaction prediction score obtained for residues 70-VVLLG-74 was 31.605, with a cutoff of 29.92 and P-value 0.112.

### Nickel Affinity Purification of SUMOylated FLAG-dyskerin

For analysis of SUMOylated FLAG-dyskerin by immunoblotting, HEK293 cells expressing 6xHis-SUMO3 and/or FLAG-dyskerin (wildtype, K446X, or SUMO3-K446X) were lysed under denaturing conditions. Briefly, cells were washed with 1XPBS and collected by scraping. One fifth of cells per condition were kept for input and lysed in 2xLaemmli followed by boiling. The remainder of the cell pellet was lysed in 6M GuHCl buffer (10mM Tris-HCl pH8, 6M GuHCl, 10mM imidazole, 0.1M NaH_2_PO_4_, adjusted to pH8 with NaOH) at room temperature by passage through a 21G1¼ syringe (5x) followed by passage through an insulin syringe (3x). Cell lysate was cleared by centrifugation at 13000rpm for 20min at 4°C. The supernatant was incubated with NiNTA resin (pre-washed 2x with 1XPBS and 1x with GuHCl buffer) on a rotator at room temperature overnight. Resin was then washed 1x with GuHCl buffer, 1x with wash buffer 1 (10mM Tris-HCl pH8, 8M urea, 10mM imidazole, 0.1M Na H_2_PO_4_, adjusted to pH8 with NaOH), and 2x with wash buffer 2 (10mM Tris-HCl pH8, 8M urea, 10mM imidazole, 0.1M NaH_2_PO_4_, 0.1% v/v Triton X-100, adjusted to pH6.3 with NaOH). For elution, resin was incubated in elution buffer (50mM NaH_2_PO_4_, 300mM NaCl, 500mM imidazole, adjusted to pH8) for 3h on a rotator at 4°C. The eluate was collected by centrifugation and resin discarded.

### Immunofluorescence

To assess localization of FLAG-dyskerin to the nucleolus, HEK293 cells expressing FLAG-dyskerin constructs were fixed with 4% formaldehyde-PBS for 10 minutes at room temperature. The fixing solution was removed and coverslips were briefly rinsed with PBS, followed by permeabilization of cells with 0.1% Triton X-100-PBS for 5 minutes at 4°C. Permeabilized cells were then washed with PBS before blocking in 5% BSA-PBS for 1 hour at room temperature. Cells were probed for FLAG-dyskerin with rabbit anti-FLAG (Sigma-Aldrich F7425, 1:500) or mouse anti-dyskerin (Santa Cruz H-3, 1:25) in PBG (1% cold fish water gelatin, 0.5% bovine serum albumin (BSA), in PBS) overnight at 4°C in a humidity chamber. In the case of assessing localization of only exogenous FLAG-tagged dyskerin, this was followed by probing with mouse anti-fibrillarin (monoclonal antibody 72B9 obtained from Dr. Kenneth Michael Pollard, 1:30) as a nucleolar marker, in PBG at 37°C for 1 hour. Coverslips were washed with PBS and immunostained in PBG with secondary antibodies conjugated to fluorescein isothiocyanate (FITC) (donkey anti-mouse IgG; Jackson ImmunoResearch Lab, Inc., 1:125) or Cy3 (donkey anti-rabbit; Jackson ImmunoResearch Lab, Inc., 1:125). Coverslips were washed with PBS and mounted in Vectashield with DAPI (Vector Laboratories). Cells with FLAG-dyskerin signal were manually scored based on localization phenotype as a percentage of the number of cells with FLAG signal detected, ≥50 cells were counted in experimental triplicate for scoring of localization of each FLAG-tagged dyskerin construct. Images were captured using an Axio Imager 2 microscope (63X; Carl Zeiss, Jena, Germany). Nucleolar localization was determined by co-localization with fibrillarin clusters, nucleoplasmic localization was determined by co-localization with DAPI, and cytoplasmic localization was determined by concentrated signal outside of and surrounding DAPI.

### Immunoprecipitation

Protein-protein interactions were assessed by immunoprecipitating FLAG-dyskerin wildtype or N/NoLS variants from HEK293 cells and immunoblotting for endogenous dyskerin-interacting proteins; by immunoprecipitating 3xFLAG-GAR1 wildtype or 5A and immunoblotting for endogenous dyskerin. Monoclonal M2 mouse anti-FLAG antibody (Sigma-Aldrich F3165) and Protein G Sepharose (GE Healthcare) pre-blocked in 1% BSA-PBS were used to immunoprecipitate (IP) FLAG-tagged and 3xFLAG-tagged proteins. The protocol used to assess protein-protein interactions was the same used to analyze the interaction between FLAG-dyskerin and hTR, and was modified based on a protocol that has been previously described for another hTR-interacting protein (88), as well as used for FLAG-tagged dyskerin (16). Briefly, cells were first lysed in low salt buffer (25mM HEPES-KCl pH7.9, 5mM KCl, 0.5mM MgCl_2_, 0.5% NP-40, 1X protease inhibitor cocktail from Roche, 20mM N-ethylmaleimide, and 40U/ml RNAseOut) for 10min on ice. Lysates were cleared by centrifugation at 5000rpm for 5min at 4°C, supernatants were kept on ice, and pellets underwent a second lysis in high salt buffer (25mM HEPES-KCl pH7.9, 350mM NaCl, 10% w/v sucrose, 0.01% NP-40, 1X protease inhibitor cocktail from Roche, 20mM N-ethylmaleimide, and 40U/ml RNAseOut) with 30sec vortex followed by 30min on a rotator at 4°C. Both low salt and high salt lysates were then cleared by centrifugation at 13000rpm for 30min at 4°C, supernatants were pooled, and total lysate was pre-cleared at 4°C on a rotator for 30min using Protein G Sepharose that was pre-washed with 1XPBS. Bradford analysis was used to calculate total protein concentration prior to IP. Lysates were incubated with anti-FLAG antibody for 2h at 4°C on a rotator before pre-blocked Protein G Sepharose was added, followed by an additional 1h incubation at 4°C on a rotator. IPs were washed 4x with 1ml of modified RIPA buffer (50mM Tris-HCl pH8, 150mM NaCl, 10mM MgCl_2_, 1% NP-40, 0.5% sodium deoxycholate, 1mM PMSF, 0.1X protease inhibitor cocktail from Rocher, and 20mM N-ethylmaleimide). For protein-protein interactions, elution from Protein G Sepharose was performed with Laemmli buffer and boiling. For protein-RNA interactions, elution was performed with TRIzol reagent (Invitrogen), followed by chloroform extraction and reverse transcription. Inputs (10% of lysate volume used for IP) were collected after pre-clearing with Protein G Sepharose and prior to IP, and treated with either Laemmli buffer and boiled, or with TRIzol reagent.

### Immunoblotting and Antibodies

Analysis of protein expression and IP experiments was performed by resolving proteins by SDS-PAGE, transfer to PVDF and immunoblotting. Primary antibodies used for immunoblotting were: anti-FLAG (Proteintech, 20543-1-AP, 1:4000) or anti-FLAG (Proteintech, 66008-3-Ig, 1:4000), anti-NAF1 (Abcam, ab157106, 1:1000), anti-NHP2 (Proteintech, 15128-1-AP, 1:5000), anti-NOP10 (Abcam, ab134902, 1:500), anti-dyskerin (Santa Cruz, sc-373956, H-3, 1:1500), anti-GAR1 (Proteintech, 11711-1-AP, 1:1000), anti-His (Santa Cruz, sc-8036, H-3, 1:500), and anti-alpha tubulin (Sigma, T5168, 1:5000).

### RNA Extraction and RT-qPCR

RNA was extracted using TRIzol reagent (Invitrogen), according to the reagent protocol. Reverse transcription was performed with SuperScript II Reverse Transcriptase (Invitrogen) according to the user protocol, with hexameric random primers. PerfeCTa SYBR Green FastMix with Low ROX (Quanta) was used for all qPCR analyses, in a 7500FAST real-time PCR system (ABI) as previously described (46). The comparative ΔΔC_T_ method was used to compare RNA enrichment between samples. For analysis of protein-RNA interactions, 5 µl of RNA from input and 5 µl of RNA from IP fractions were reverse transcribed into cDNA and subjected to qPCR using specific primers for target RNAs (**Table S1**). The ΔΔC_T_ was calculated between the mean C_T_ of the IP and the mean C_T_ of the input for each sample.

### Q-TRAP

Quantitative analysis of telomerase activity was done using the Q-TRAP protocol previously described (89). Briefly, HEK293 cells with or without expression of FLAG-dyskerin constructs were treated with scramble siRNA or siRNA to deplete endogenous dyskerin for 72h prior to harvesting by scraping and lysis in NP-40 lysis buffer. A standard curve was generated with a serial dilution of mock lysate (HEK293 cells untreated with siRNA and not expressing FLAG-dyskerin) for each experimental replicate (n=3), with 1 µg, 0.2 µg, 0.04 µg, 0.008 µg, and 0.0016 µg of total protein. For comparison of telomerase activity between conditions, 0.2 µg of total protein was used for each sample.

### Statistical Analyses

All statistical analyses were performed using GraphPad Prism 7. One-way ANOVA tests (*p* < 0.01) were used to compare RNA enrichment when assessing interaction between FLAG-dyskerin and hTR, and for comparison of relative telomerase activity (RTA) in Q-TRAP experiments. For analysis of RNA interaction, the enrichment of hTR in each N/NoLS variant IP fraction was compared to the enrichment of hTR in the FLAG-dyskerin wildtype IP fraction. For analysis of telomerase activity, the RTA percentage of each condition was separately compared to the mock (untreated HEK293 cells) RTA percentage. Each experiment was performed in triplicate, and error bars represent the standard error of the mean between experimental replicates. Dunnet’s test was used to correct for multiple comparisons.

## Acknowledgements

We thank Dr. Frédérick Antoine Mallette, Dr. François Dragon, and Dr. Ling-Ling Chen for providing us with pcDNA3.1-6xHis-SUMO3, pcDNA3.1-FLAG-dyskerin^WT^, and pmEGFP-C1-DKC1 plasmids, respectively. We thank Dr. Kenneth Michael Pollard for providing us with mouse anti-fibrillarin antibody for IF. We thank Dr. Stéphane Richard for use of the Axio Imager 2 microscope.

## Author Contributions

Author contributions: D.E. M., and C. A. designed research; D.E. M, P. L.-L., J. Q., F. M., and E. B. performed experiments; D.E. M. and C. A. wrote the manuscript; all authors contributed to reviewing and editing the manuscript.

**Declaration of Interests**

